# Can *Drosophila melanogaster* tell who’s who?

**DOI:** 10.1101/342857

**Authors:** Jonathan Schneider, Nihal Murali, Graham Taylor, Joel Levine

## Abstract

*Drosophila melanogaster* are known to live in a social but cryptic world of touch and odours, but the extent to which they can perceive and integrate visual information is a hotly debated topic. Some researchers fixate on the limited resolution of *D. melanogaster’s* optics, other’s on their seemingly identical appearance; yet there is evidence of individual recognition and surprising visual learning in flies. Here, we apply machine learning and show that individual *D. melanogaster* are visually distinct. We also use the striking similarity of *Drosophila’s* visual system to current convolutional neural networks to theoretically investigate *D. melanogaster’s* capacity for visual understanding. We find that, despite their limited optical resolution, *D. melanogaster’s* neuronal architecture has the capability to extract and encode a rich feature set that allows flies to re-identify individual conspecifics with surprising accuracy. These experiments provide a proof of principle that *Drosophila* inhabit in a much more complex visual world than previously appreciated.

**Author summary:** In this paper, we determine a proof of principle for inter-individual recognition in two parts; is there enough information contained in low resolution pictures for inter-fly discrimination, and if so does *Drosophila’s* visual system have enough capacity to use it. We show that the information contained in a 29×29 pixel image (number of ommatidia in a fly eye) is sufficient to achieve 94% accuracy in fly re-identification. Further, we show that the fly eye has the theoretical capacity to identify another fly with about 75% accuracy. Although it is unlikely that flies use the exact algorithm we tested, our results show that, in principle, flies may be using visual perception in ways that are not usually appreciated.

## Introduction

There is an increasing body of evidence that *Drosophila melanogaster* lives in a surprisingly rich and complex world, including group behaviour [1], communal learning [2], and recognition during aggressive behaviours [3]. This repertoire of social behaviour has often been assumed to be independent of vision, as *Drosophila’s* compound eye was thought to have insufficient visual acuity to play a serious role. With only ^~^850 lens units (ommatidia), each capturing a single point in space, *Drosophila’s* compound eye resolution is certainly low. And the level of detail, traditionally determined by the inter-ommatidial angle, renders anything but movement or regular patterns seemingly impossible to discern (Fig. 1B).

**Fig 1.**
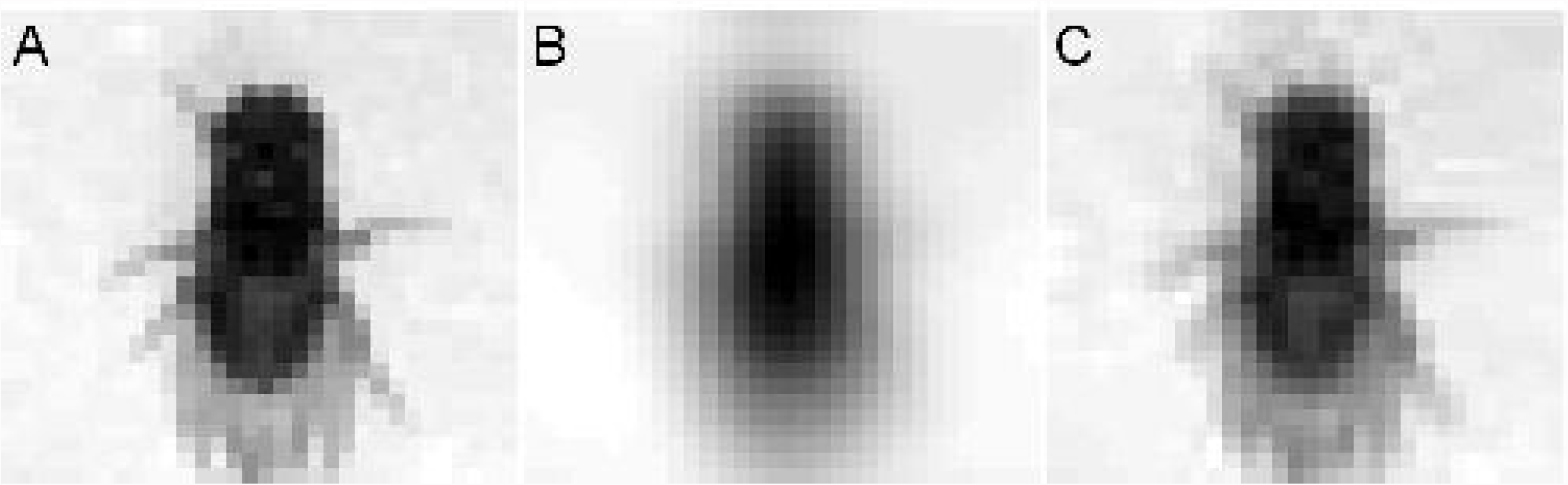
Theoretical visual acuity of *Drosophila melanogaster.* Image of *Drosophila melanogaster* represented after various theoretical bottlenecks. A: Image of a female *D. melanogaster* re-sized through a 32×32 bottleneck. B: The same image, but adjusted using AcuityView [4] for a viewing distance of ^~^3 body lengths using the inter-ommatidial angle of 4.8° [5]. C: The same image and distance, but using a conservative estimate of the effective acuity determined by Juusola *et al.* [6] of ^~^1.5°.

However, recent physiological experiments [6] reveal that *D. melanogaster* can respond to details as fine as 1.16°as long as they are presented (to a tethered fly) at specific speeds. These speeds happen to coincide with *D. melanogaster’s* natural saccadic gait [7], which strongly suggests that naturally behaving *D. melanogaster* have much finer than the inter-ommatidial angle of 4.8° [5]. This hyper-acuity is found at the photoreceptor level (due to rhabdomere movement changing the angle of light reception), allowing most of the capacity of the visual network to be devoted to information processing. At this effective hyper-acuity, and at socially relevant distances, the number of ommatidia and not the inter-ommatidial angle becomes the limiting factor (Fig. 1). This acuity potentially puts them in the same visual league (albeit with lower resolution) as *Apis mellifera* [8], which has been able to, among other visual feats, identify individual human faces [9].

This spatio-temporal coding and increased visual acuity potentially explains recent studies which have shown that *D. melanogaster* can not only resolve other flies, but can also decode social meaning using vision (e.g. female choice of male phenotypes [10] and exposure to parasitoids [11]). Combined, these results open up the possibility that *Drosophila* uses vision to a much greater extent in object recognition, perhaps even using it to discriminate between species or sex (supplementing other olfactory cues that are known to convey this information [12]).

Even with *D. melanogaster’s* photoreceptor hyper-acuity [6], the image that is received is only around 29×29 units (or pixels; Fig. 1). We wanted to know if there is enough absolute information contained in this low-resolution image to identify individuals from each other. One approach is to task Deep Convolutional Networks (DCNs) to differentiate individual *D. melanogaster,* as DCNs are engineered to learn/extract/use any useful features found in the images. If there is enough individual-level variation for highly engineered DCNs, we would want to investigate the possibility that *D. melanogaster* also take this low resolution image and extract meaningful information out of it. Should individual flies prove visually unique and *D. melanogaster’s* visual network have enough capacity, vision could potentially play a role in identifying beyond species or sex, perhaps aiding in determining familiar or non-familiar conspecifics in social situations [3].

*How* a fly’s visual system could extract meaning out of low resolution images is suggested by the highly structured and layered organization of *Drosophila’s* visual system (Fig. 2C). At the input the ommatidia are packed one by one, but their individually-tuned photoreceptors are arranged spatially to essentially convolve a 6-unit filter across the receptive field. The output of this photoreceptor filter is, in turn, the input for downstream medulla neurons that connect to several ‘columns’ of photoreceptor outputs. This filter-convolution and using the output of one filter as a ‘feature map’ for another layer is a hallmark of the engineered architectures of DCNs that dominate computer vision today (one such DCN is illustrated in Fig. 2A). Just as DCNs can take low level image representations and encode them into a semantic representation, *D. melanogaster’s* visual system seems well-suited to similarly build up semantic meaning in images.

**Fig 2.**
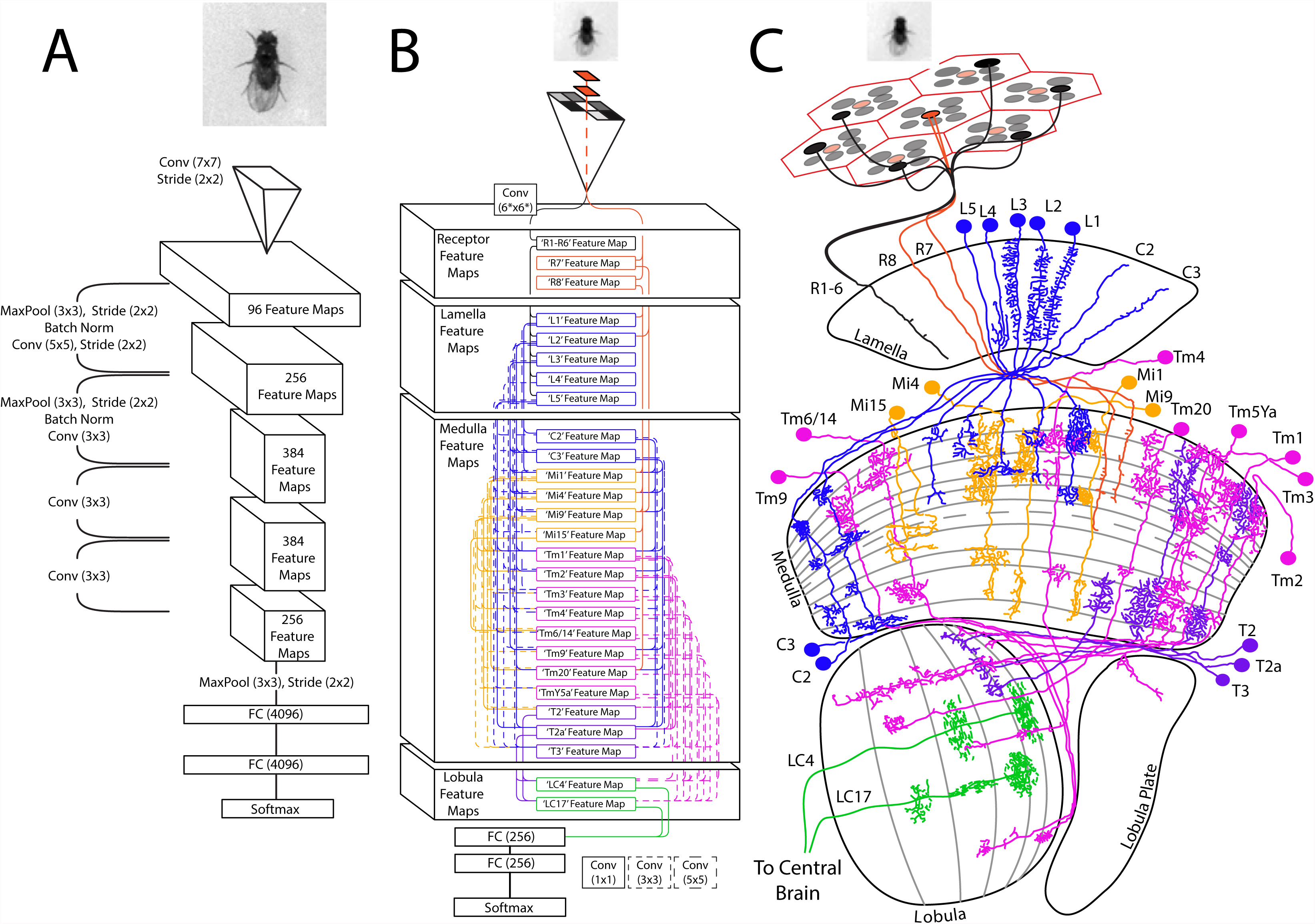
Our fly-eye merges engineered and biological architectures. Schematics of a ‘standard’ convolutional network, our fly-eye model, and a simplified visual connectome of *Drosophila.* A: Architecture of Zeiler and Fergus [13], receiving the original 181×181 pixel image of an individual *Drosophila melanogaster.* B: Our fly-eye model, receiving a 29×29 scaled-down image of an individual *Drosophila,* and showing connections between feature maps. Besides the first custom 6-pixel convolutional filter and the 1×1 convolutional filters (‘R7’ and ‘R8’), all other convolutions are locally connected filters. C: A simplified map of the fly visual circuit receiving the same scaled-down image of another *D. melanogaster.* The connections among the neurons implemented in our model are displayed, illustrating the connections and links within and between layers (adapted from [14] and [15]). See S2 Table for performance of these models on a traditional image-classification dataset.

We investigate whether *D. melanogaster* could theoretically categorize and recognize its complex visual environment. To determine how much absolute underlying visual variation is available for social behaviour in *D. melanogaster,* we examine the ability of humans and human-inspired deep convolutional models to re-identify individual *D. melanogaster* across days. To see whether *D. melanogaster* is *capable* of using this individual-level visual variation between flies, we investigate a model of *Drosophila’s* visual system in a conspecific re-identification paradigm. This study builds upon the behavioural results of conspecific information and physiological evidence of hyper-acute vision, and provides a proof of principle to dispel the oft-touted argument that *D. melanogaster’s* visual ability is limited to low-level object- and pattern-detection. Here we present evidence that *D. melanogaster* may likely see and live in a much richer social environment than is appreciated.

## Materials and methods

### Simplified *Drosophila* model eye

We implemented a virtual fly visual system using standard deep learning libraries (Keras; code available at github). Our implementation uses ^~^25,000 artificial neurons whereas *Drosophila* has ^~^60,000 neurons in each visual hemisphere [16]. We purposefully did not model neurons that are structurally suggestive to respond to movement, and therefore we were specifically limited to ‘modular’ neurons (with 1 neuron/column) throughout the medulla [17]. The connections between neuronal types was extracted from published connectomes [18]. We imposed artificial hierarchy on our model eliminating self connections between neuron ‘subtypes’ (i.e. no connections between L1 and L1, or L1 and L2), and while we allowed initial layers to feed into multiple downstream layers, we eliminated ‘upstream’ connections. The final lobula-like artificial neurons were modelled after [15]. The layers were ordered according to their axon penetration deeper into the system. The model is illustrated in Fig. 2B, beside the biological inspiration (Fig. 2C). See S1 Table for complete connection map and hierarchy, and S2 Table for comparative performance of this models on a traditional image-classification dataset.

### Fly Data Acquisition

*D. melanogaster* were housed at 25C°on a 12-12 Light-Dark cycle. 10 males and 10 females were collected 1-4 hours after eclosion and housed separately. On the third day post-eclosion flies were individually mouth pipetted into a circular acrylic arena (60mm diameter, 2mm high). These flies were illuminated with standard overhead LED bulbs and filmed in grayscale with a GRAS-20S4M for 15 minutes, with 16 frames/second. This was repeated for three consecutive days in total, which resulted in 14,400 ×3 images per fly. Each filming session was within 2 hours of ZT 8. Three independent datasets were acquired of 20 flies each.

### Fly Data Processing

Each video was tracked using CTRAX [19], and the tracked position and orientation was used to localize the flies and orient the images so that the flies were always centred and facing ‘up’. The training set for each week was constructed from days 1 and 2 equally, and consisted of the first 75% of the each fly’s recording (12240 frames). The validation set was the final 15% of the recording (2160 frames). The test dataset was the entire recording on day 3. Images were standardized by subtracting the mean and dividing by the standard deviation of the training set. For ResNet18 [20] and the Zeiler and Fergus [13] models, the original 181×181 image was either: reduced to 33×33 then centre-cropped to 29×29 and then re-sized to 224×224 or re-sized to 256×256 and centre-cropped to 224×224 (effectively using the centre 158×158 pixels).

### Human Performance

A GUI program was written in MATLAB which presented a human observer with 3 viewpoints of an exemplar fly: dorsal, ventral, and sideways obtained from the first two days of filming. The observer was then asked to choose among 20 images (of the 20 flies) obtained from day 3, of which one belong to the exemplar (S3 Fig-S4 Fig). Note that this is a compare/match setup rather than a learn/recall. The pictures were randomly re-sized through a 29×29 bottleneck.

## Results

We also wanted to see whether various architectures, both biologically rooted and not, could detect visual differences between flies across days (a notably non-human task). We acquired three rounds of images, each round having 10 males and 10 females, filmed for 3 consecutive days. Knowing that age/experience may slightly affect the morphology of flies, we experimented with training these networks on days 1 and 2, and testing their ability to re-identify the flies on day 3. We evaluated the efficient and high-capacity model (ResNet18 [20]), a model that rivals human representation (Zeiler and Fergus [13]), our fly-eye model, and human performance. These results are summarized in Table 1.

**Table 1.** Performance On *D. melanogaster* Re-Identification

| Model Name | Resolution<sup>1</sup><br>(pixels) | Accuracy<br>(F<sub>1</sub> Score)<sup>2</sup> |
| --- | --- | --- |
| ResNet18 [20] | 158×158 | 0.9426 ±0.0358 |
| Zeiler and Fergus [13] | 158×158 | 0.9373 ±0.0365 |
| Human Performance | 158×158 | 0.1309 |
| Zeiler and Fergus [13] | 29×29 | 0.8549 ±0.0778 |
| ResNet18 [20] | 29×29 | 0.8357 ±0.0909 |
| **Our fly-eye** | 29×29 | **0.7548 ±0.1141** |
| **Our fly-eye**<br>**w/ random zoom<sup>3</sup>** | 29×29 | **0.5486 ±0.1316** |
| Human Performance | 29×29 | 0.0829 |
| Random Chance |  | 0.05 |

Mean and standard deviation shown (n=3 independent datasets)
<sup>1</sup> For Zeiler and Fergus and ResNet18, the “resolution” was a bottleneck (see methods).
<sup>2</sup> An example of high precision / low recall can be seen in S2 Table as ID 10 is assigned to the right fly 94% of the time, but fly 10 is only correctly identified 37% of the time.
<sup>3</sup> The zoom was applied randomly without preserving aspect ratio (see S2 Fig).

As a benchmark, we applied an ResNet18 architecture (see S1 Fig). This was our highest performing network architecture, achieving an F_1_-score of 0.94 (with high resolution over the three datasets). While the average performance is good, we note certain problematic individual flies that become difficult to accurately re-identify across days (e.g. in week 2, fly 10’s accuracy on day 3 was 37% with equal confusion between two other flies S2 Table). Forcing the images through a bottleneck (which ensures that the information content is similar to the reduced-resolution fly-eye model) decreased the Fl-score by 0.11 for ResNet18. The Zeiler and Fergus [13] architecture was the most robust in terms of the bottleneck, only decreasing the F_1_-score by 0.08, but did not achieve as high a performance as ResNet18.

The fly-eye model achieves a relatively high F_1_ score of 0.75, which is not that much lower than the highly engineered ResNet18 (at low resolution). To eliminate the fly-eye model’s ability to measure absolute size/shape and force relative feature extraction, we randomly re-sized the images (both training and testing) by up to 25% without preserving proportion (see S2 Fig for examples). Our virtual fly-eye outperformed humans, even without absolute size measures, achieving an F_1_-score of ^~^0.55 re-identifying conspecifics. We also note that the fly-eye model almost never mistakes a male for a female (F_1_-score exceeding 0.99 when the re-identification IDs were collapsed by sex).

To establish a human-performance baseline we used volunteers to attempt the re-identification of flies (S3 Fig-S4 Fig). This is an especially challenging task given the range of motion and viewpoints that an individual fly has in an unconstrained arena. As this task falls outside normal visual object recognition, we restricted volunteers to be very experienced “fly-pushing” scientists. Human performance was predictably low, but varied, with an average Fl-score of 0.11 (.08 when the images were reduced to 29×29 pixels, 0.13 when given the full resolution 181×181 images).

## Discussion

Our results suggest that *Drosophila* have the innate capacity to extract semantic meaning from their visual surrounding, and even if we are still discovering *how Drosophila* could encode its world, we should not disregard its visual understanding.

We note that our fly-eye model performs poorly on a standard image classification task (F_1_ Score of ^~^0.40 on CIFAR10; see S2 Table) which has built in ranges of zoom and positional variance of the objects. One possible explanation of the inability to cope with different object sizes is that, unlike other architectures, *D. melanogaster’s* visual system maintains the input dimensions (the columnar medulla neurons). DCN’s dimension reduction (and other ‘tricks’ like pooling layers) convey a larger position-invariance to low level feature detectors. Without them, our fly-eye model only does well when the distance to the object is fixed. It is tempting to posit that an individual can therefore have an innate and/or experience-dependent distance at which visual information is preferentially understood, and that this could be one of the determinants for social spacing and interaction distances [21, 22].

One prediction that arises from this model and its apparent ability to encode more than simple “looming” and “movement” visual cues is that the highest (furthest from input) level feature maps may correspond to semantically rich meaning in the visual system. These lobula neurons should then encode (among other information) complex object recognition categories and stimulating them should produce more than simple object-avoidance behaviours in *Drosophila.* While some lobula columnar neurons (like LC11) seem specialized for high-acuity small object motion detection [23], others appear to encode more complex information. These other LC neurons (like LC17), when stimulated, seem to provoke social-context dependent behaviours [15].

We are aware of other investigations using DCNs to classify insect species [24]. However the most relevant study which involve organism re-identification only does so over short (<1 min) time frames (IDTracker2.0 [25]). Prior to this study, DCNs have only been effective on images that are temporally very close [25]. We observe a non-general loss of accuracy for specific flies, with some flies having an accuracy lower than 40% (S2 Table). This ability to re-identify flies *across* days opens experimental possibilities, especially considering that this performance was achieved with static images (16fps yields around a thousand estimates of ID per minute, allowing high confidence in the parsimonious correct identification). This is in contrast to the human ability to re-identify flies, which at low resolutions is barely better than chance.

Clearly, all models can learn to re-identify flies to some extent, underscoring the individual-level variation in *D. melanogaster.* Re-identifying flies is in fact easier for DCNs than CIFAR10 (at least with centred images of flies acquired at the same distance). Even the model that rivals, in some sense, the representational performance of humans [26] does *ten times* better than humans. Why humans can’t tell one fly from another is not clear, regardless of whether it was evolutionarily beneficial to discriminate individual flies, humans do have incredible pattern detection abilities. It may simply be a lack of experience (although we attempted to address this by only using experienced *Drosophila* researchers as volunteers) or a more cryptic pattern-recognition ‘blind-spot’ of humans. In either case, these findings should spur new experiments to further understand the mechanisms of human vision and experience and how they fail in this case.

Machine learning practitioners are pushing for deep networks to use more biologically-inspired design choices and training algorithms [27–29]. Neurobiologists could use these models to generate hypotheses in how information is processed in the visual system. We think *D. melanogaster* is well-suited to link these two fields to continue unravelling evolution’s solution to visual processing. This new area offers a simple, genetically- and experimentally-tractable organism to peer into the workings of the visual system which will no doubt uncover not just how *Drosophila,* but all of us, see the world.

## Conclusion

These results help explain recent, traditionally controversial, findings that *Drosophila melanogaster* can resolve relatively detailed visual semantic meaning (e.g. female choice of males [2] and parasitoids exposure [11]). We show here that each *D. melanogaster* has visually distinguishable features that persist across days. This fact, combined with their hyper acuity [6] and theoretical capacity of their visual network, provides a proof of principle against the traditional belief that *Drosophila* only see blurry motion. In fact, *Drosophila* could have the ability to see and discriminate a surprisingly diverse visual world, in some cases possibly better than we can.

## Supporting information

**S1 Fig. ResNet18 Architecture.** Residual Networks with 18 ‘layers’ [20] were constructed as depicted, using the improved block scheme proposed by [20] (top left inset).

**S2 Fig. Examples of fly pictures.** On the left are examples of the various viewpoints of the fly. The pictures indicated were run through the random non-proportion-preserving zoom to generate three examples of each.

**S3 Fig. Human Query: High resolution example.** Participants were asked to re-identify the exemplar fly (top left) which exhibited three viewpoints: ventral, dorsal, and angled. They could request new pictures of said fly as many times as needed. The exemplar pictures were manually sorted into viewpoint categories and then randomly selected from Days 1 and 2. The other fly pictures (queries) were selected from Day 3.

**S4 Fig. Human Query: Low resolution example.** Participants were asked to re-identify the exemplar fly (top left) which exhibited three viewpoints: ventral, dorsal, and angled. They could request new pictures of said fly as many times as needed. The exemplar pictures were manually sorted into viewpoint categories and then randomly selected from Days 1 and 2. The other fly pictures (queries) were selected from Day 3.

**S1 Appendix. Connectome of the model fly-eye**

Constructed from the published connectome [18]. We imposed a hierarchy (see text), but otherwise allowed links between ‘lower’ layers as long as at the links were reported at least once (orange). Links between layers and connecting ‘higher’ levels were not used (blue). In brief, a 6-pixel filter is convolved through the image (representing photoreceptors R1-R6; whether or not the image is grayscale, the filter is fixed in all channels) and two additional colour-sensing filters are convolved (representing R7/R8; 1×1 pixel filter). The output of R1-R6 are then used as the feature map for lamina neurons L1-L5, which are locally connected 1×1 filters (i.e. different filters are learned at each spatial position). The outputs of these L1-L5s locally convolved and are fed into the medulla intrinsic (Mi) neurons and/or the centrifugal (C) neurons, and/or the transmedullary (Tm) neurons. The C neurons feed into the Mi, Tm, T neurons. The Mi neurons feed into the Tm, and T neurons. The Tm neurons apply a filter and send their outputs to the T neurons. Sizes of the filters were determined from Takemura *et al.* [18], who traced connections through a single focal column (labelled Home) and up to two columns in any direction (in a hex grid they are labelled A-R). If a previous column had only connections to its respective column (i.e. Home→Home, A→A), it was modelled with a 1 × 1 locally-connected filter. If a previous column had more than 3 connections to its immediate surrounding columns (i.e. Home→A, C→D) then it was modelled with a 3×3 locally-connected filter. Finally, if it had more than 3 connections to more distant neighbours (i.e. Home→J, P→A), then it was modelled with a 5×5 locally-connected filter. Unfortunately the connections between the medulla, lobula and brain are not as documented as those between and within the lamella and medulla but we implement a lobula neuron-like LC17 that concatenates Tm and T neurons with a 3×3 filter, while another neuron (LC4-like) concatenates Tm and T neurons with a 5×5 filter [15]. The output feature maps are then flattened and fed into two densely connected layers with 256 neurons each before a soft-max layer.

**S1 Table.**
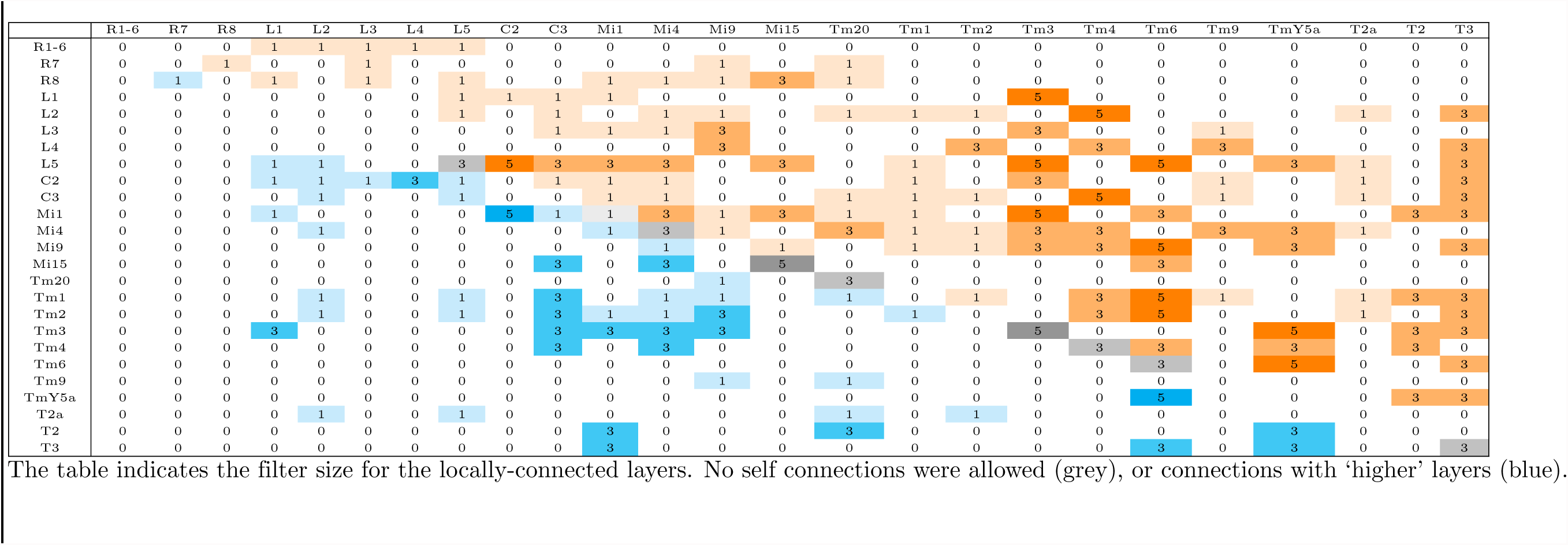
Model fly-eye connectome.

## CIFAR10

### CIFAR10 Data Processing

Images were standardized by subtracting the mean and dividing by the standard deviation of the training set for each colour channel. In all cases, to be comparable, the images were processed to trim a row and column from the top and left, and two rows and columns from the bottom and right (trimming to 29×29 pixels resulted in higher accuracy than resizing from 32×32) and were minimally augmented (random vertical flips and each image randomly offset by 3 pixels). For ResNet18 [20] and the Zeiler and Fergus [13] models, images were re-sized to 224×224.

### CIFAR10 Results

The CIFAR-10 dataset consists of colour images (32×32 pixels) in 10 classes (airplane, automobile, bird, cat, deer, dog, frog, horse, ship and truck) [30]. The current state of the art models can achieve 97.44% accuracy (with clever data augmentations [31]), while human performance has been estimated at around 94% accuracy [32]. Our re-implementation of ResNet18 [20] achieves 0.91 (F_l_ score). The Zeiler-Fergus model [13], that has been shown to rival the representational performance of the human inferior temporal cortex [26] (Illustrated in Fig. 2A), achieves a lower Fl score of 0.85, revealing the gap between the ability to represent mid-level complexity and highest order syntactic information. Our simplified implementation of the fly visual system achieves 0.40. The CIFAR10 results are summarized in S1 Table.

**S2 Table Results of a simple vision task (CIFAR10).**

**Table 2.** Performance On CIFAR10

| Model Name | ‘Neurons’<br>(#) | Parameters<br>(#) | Accuracy<br>($F_1$ Score)<sup>1</sup> |
| --- | --- | --- | --- |
| Human Performance | Billions |  | 0.94<sup>2</sup> |
| ResNet18 [20] | ~2 million | ~11 million | 0.9148<sup>3</sup> |
| Zeiler and Fergus [13] | ~1.6 million | ~72 million | 0.8488<sup>3</sup> |
| **Our fly-eye** | 25,742 | 1,350,316 | **0.5364** |
| Random Chance |  |  | 0.10 |

<sup>1</sup> The $F_1$ score combines precision (probability of assigning the right ID to the right class) and recall (probability that the ID assigned is to the right class).
<sup>2</sup> Human performance is given in overall accuracy (%).
<sup>3</sup> Images were re-sized from $29 \times 29$ to $224 \times 224$ . Their results presented here are not for state-of-the-art benchmarking, but for comparison.

**S2 Table.** Confusion Matrix for ResNet18 Week 2.

|  | 1 | 2 | 7 | 3 | 10 | 9 | 6 | 8 | 4 | 5 | 13 | 14 | 20 | 18 | 16 | 17 | 15 | 11 | 19 | 12 |
| --- | --- | --- | --- | --- | --- | --- | --- | --- | --- | --- | --- | --- | --- | --- | --- | --- | --- | --- | --- | --- |
| 1 | 93 | 0 | 1 | 0 | 0 | 3 | 1 | 1 | 1 | 0 | 0 | 0 | 0 | 0 | 0 | 0 | 0 | 0 | 0 | 0 |
| 2 | 0 | 98 | 1 | 0 | 0 | 0 | 0 | 0 | 0 | 0 | 0 | 0 | 0 | 0 | 0 | 0 | 0 | 0 | 0 | 0 |
| 7 | 0 | 8 | 89 | 0 | 1 | 2 | 0 | 0 | 0 | 0 | 0 | 0 | 0 | 0 | 0 | 0 | 0 | 0 | 0 | 0 |
| 3 | 0 | 0 | 0 | 99 | 0 | 0 | 0 | 0 | 0 | 0 | 0 | 0 | 0 | 0 | 0 | 0 | 0 | 0 | 0 | 0 |
| 10 | 0 | 29 | 27 | 1 | 37 | 1 | 0 | 1 | 2 | 1 | 0 | 0 | 0 | 0 | 0 | 0 | 0 | 0 | 0 | 0 |
| 9 | 0 | 0 | 0 | 0 | 0 | 99 | 0 | 0 | 0 | 0 | 0 | 0 | 0 | 0 | 0 | 0 | 0 | 0 | 0 | 0 |
| 6 | 0 | 1 | 0 | 0 | 0 | 0 | 98 | 0 | 1 | 0 | 0 | 0 | 0 | 0 | 0 | 0 | 0 | 0 | 0 | 0 |
| 8 | 0 | 0 | 0 | 0 | 0 | 0 | 0 | 99 | 0 | 0 | 0 | 0 | 0 | 0 | 0 | 0 | 0 | 0 | 0 | 0 |
| 4 | 0 | 0 | 0 | 0 | 0 | 0 | 0 | 0 | 99 | 0 | 0 | 0 | 0 | 0 | 0 | 0 | 0 | 0 | 0 | 0 |
| 5 | 1 | 1 | 2 | 0 | 0 | 3 | 2 | 3 | 4 | 82 | 0 | 0 | 0 | 0 | 0 | 0 | 0 | 0 | 0 | 0 |
| 13 | 0 | 0 | 0 | 0 | 0 | 0 | 0 | 0 | 0 | 0 | 95 | 1 | 1 | 2 | 0 | 2 | 0 | 0 | 0 | 0 |
| 14 | 0 | 0 | 0 | 0 | 0 | 0 | 0 | 0 | 0 | 0 | 1 | 98 | 0 | 1 | 0 | 0 | 0 | 0 | 0 | 0 |
| 20 | 0 | 0 | 0 | 0 | 0 | 0 | 0 | 0 | 0 | 0 | 0 | 0 | 97 | 0 | 0 | 1 | 0 | 0 | 0 | 0 |
| 18 | 0 | 0 | 0 | 0 | 0 | 0 | 0 | 0 | 0 | 0 | 0 | 0 | 0 | 97 | 0 | 1 | 0 | 0 | 0 | 2 |
| 16 | 0 | 0 | 0 | 0 | 0 | 0 | 0 | 0 | 0 | 0 | 0 | 0 | 0 | 2 | 70 | 0 | 0 | 4 | 0 | 24 |
| 17 | 0 | 0 | 0 | 0 | 0 | 0 | 0 | 0 | 0 | 0 | 0 | 0 | 0 | 3 | 0 | 96 | 0 | 0 | 0 | 0 |
| 15 | 0 | 0 | 0 | 0 | 0 | 0 | 0 | 0 | 0 | 0 | 0 | 0 | 0 | 5 | 0 | 0 | 93 | 0 | 0 | 1 |
| 11 | 0 | 0 | 0 | 0 | 0 | 0 | 0 | 0 | 0 | 0 | 0 | 0 | 0 | 0 | 0 | 0 | 0 | 99 | 0 | 1 |
| 19 | 0 | 0 | 0 | 0 | 0 | 0 | 0 | 0 | 0 | 0 | 1 | 6 | 0 | 1 | 1 | 7 | 0 | 1 | 82 | 2 |
| 12 | 0 | 0 | 0 | 0 | 0 | 0 | 0 | 0 | 0 | 0 | 0 | 0 | 0 | 1 | 0 | 0 | 0 | 0 | 0 | 98 |

Note that fly 10 has a recall accuracy of 37%, and is almost equally confused for flies 2 and 7. Flies are ordered by sex (Purple = male, Yellow = Female), then by ascending size. Predictions are colour coded and weighted by percentage (correct predictions are indicated in orange, incorrect predictions are coloured cyan).

## Acknowledgments

We thank Becci Rooke for comments and suggestions and the following volunteers who were tasked with fly-reidentification: Jacob Jezovitz, Mireille Golemiec, Amara Rasool, and Nawar Awash.

